# Literature Consistency of Bioinformatics Sequence Databases is Effective for Assessing Record Quality

**DOI:** 10.1101/101873

**Authors:** Mohamed Reda Bouadjenek, Karin Verspoor, Justin Zobel

## Abstract

Bioinformatics sequence databases such as Genbank or UniProt contain hundreds of millions of records of genomic data. These records are derived from direct submissions from individual laboratories, as well as from bulk submissions from large-scale sequencing centres; their diversity and scale means that they suffer from a range of data quality issues including errors, discrepancies, redundancies, ambiguities, incompleteness, and inconsistencies with the published literature. In this work, we seek to investigate and analyze the data quality of sequence databases from the perspective of a curator, who must detect anomalous and suspicious records.

Specifically, we emphasize the detection of inconsistent records with respect to the literature. Focusing on GenBank, we propose a set of 24 quality indicators, which are based on treating a record as a query into the published literature, and then use query quality predictors. We then carry out an analysis that shows that the proposed quality indicators and the quality of the records have a mutual relationship, in which one depends on the other. We propose to represent record-literature consistency as a vector of these quality indicators. By reducing the dimensionality of this representation for visualization purposes using Principal Component Analysis, we show that records which have been reported as inconsistent with the literature fall roughly in the same area, and therefore share similar characteristics. By manually analyzing records not previously known to be erroneous that fall in the same area than records know to be inconsistent, we show that 1 record out of 4 is inconsistent with respect to the literature. This high density of inconsistent record opens the way towards the development of automatic methods for the detection of faulty records. We conclude that literature inconsistency is a meaningful strategy for identifying suspicious records.

## 1 Introduction

Bioinformatics sequence databases such as Genbank or UniProt contain large numbers of nucleic acid sequences and protein sequences. In 2016, GenBank^1^ alone contained over 224 billion nucleotide bases in more than 198 million sequences – a number that is growing at an exponential rate, doubling every 18 months.^2^ In commercial organizations, the primary reason for creating and maintaining such databases is their importance in the process of drug discovery [1]. Thus a high level of data quality is crucial.

However, since these databases are fed by direct submissions from individual laboratories and by bulk submissions from large-scale sequencing centers, they suffer from a range of data quality issues [2] including errors, discrepancies, redundancies, ambiguities, and incompleteness. In this work, we seek to investigate and analyze the data quality of sequence databases from the perspective of a curator, who must detect anomalous and suspicious records.

Usually, a suspicious record is reported manually, by one of a curator whose job consists mainly of checking the database records, the record’s original submitter, or a third person who is a user of the database and has noticed the inconsistency of that record. To illustrate this difficult job of spotting erroneous records, in Figure 1, we show the distribution of record ages using boxplot, which is a convenient way of graphically depicting groups of numerical data through their quartiles. Hence, Figure 1 shows the distribution of the ages of records that are marked as “alive” (active) and “dead” (removed); note that this is the terminology used by curators. The age of an “alive” record is computed based on the date on which this paper has been written, e.g., for a record that has been created exactly one year ago, its age is 365 days in the distribution presented in Figure 1. However, the age of a “dead” record is computed based on its removal date, e.g., if a record is created and then removed after 10 days of existence, its age is obviously 10 days in the distribution presented in Figure 1. Figure 1 leads us to make two hypotheses: since removed records have an average age of about 1 month at their removal time, either (i) it takes about one month for a problematic record to be detected, or (ii) curators focus only on new records, while neglecting older ones. Either way, it is clear that there is a time window of only 1 month during which curators act. Hence, if a suspicious record is not identified in this time window, it has a low probability of being spotted. These observations show the difficulty of the curator’s job, and the need for the development of automatic methods to assist them.

**Figure 1:**
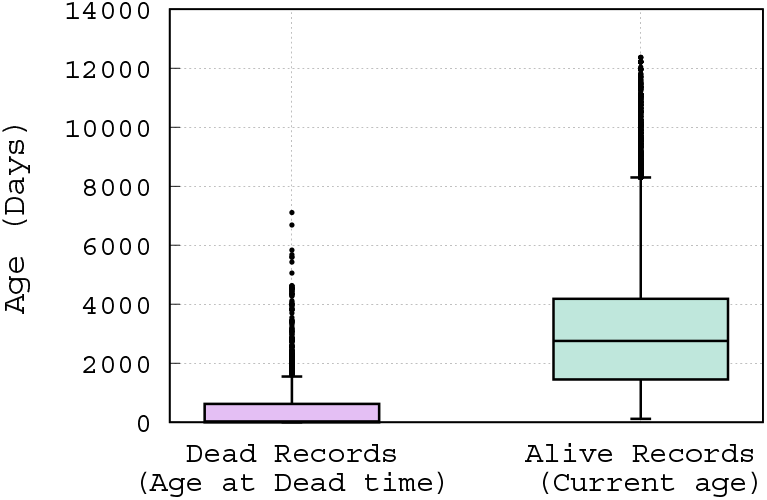
Box-plot of the distribution of record ages.

In contrast to previous research, which primarily focused on detection of duplicate records [3, 4] and erroneous annotations [5, 6], we emphasize the detection of inconsistent records with respect to the literature. In this work, we define an inconsistent record as a record that doesn’t match the content of its associated research articles. Each record cites a list of publications by the authors of the sequence that discuss the data reported in that record. However, an inconsistent record may: (i) specify a different organism than the one discussed in its associated articles, (ii) specify a different gene name (or protein name) than the one discussed in its associated articles, or (iii) focus on a gene (or a protein) that is not reported in its associated articles. For example, the record with the accession number BK004887^3^ specifies the organism “Gorilla gorilla”, whereas the article PMC555591^4^ that is supposed to report on this sequence refers to the organism “Chimpanzee”. This inconsistency has been correctly reported and this record has been removed from the database. On the other hand, the record FJ824848^5^ that describes the complete sequence of a cloning vector pDMK3^6^, has an associated article PMC2675058^7^ that describes this cloning vector as pDMK2. This record has not been reported as inconsistent, and is still alive in the database. Clearly this record should be flagged as suspicious, for review by GenBank curators who should recommend to its submitter that the vector name be corrected.

In this work, we propose that the literature linked to Genbank records in their “*REFERENCE*” fields can be used as background knowledge to check their quality. We explore a combination of information retrieval (IR) and machine learning techniques to identify records that are anomalous and thus merit analysis by a curator. Specifically, we propose a set of literature based quality features, which are used to analyze and compare records that are “alive” and other records that are “dead” based on several removal reasons. The analysis we carried out shows clearly that anomalous records removed because *“evidence is not reported/published in the related literature”*, group together in a 2D representation of the data, indicating that they have shared characteristics based on our proposed features.

In prior work, we have considered the use of the literature for data quality assessment of bioinformatics sequence databases in a supervised classification setting [7]. In that work, we demonstrated that, although fewer than 0.25% of the records in our data set are known to be faulty, by automated comparison with literature, faulty records could be identified with a precision of up to 10% and a recall of up to 30%, while strongly outperforming several baselines. However, our error analysis revealed substantial limitations of the methodology, due to the extreme imbalance of the data and the inadequacies of the available labels. This work both motivated the need for exploration of unsupervised strategies for detecting faulty records, and gave us confidence in exploring further the value of the literature for this purpose.

The contributions of this paper are as follows:

- We demonstrate that the research literature can be used in an automatic, unsupervised setting for assessing the quality of a record.
- We propose a list of quality indicators, which are based on the published literature.
- We propose a list of quality indicators for the unsupervised quality analysis, which are based on the published literature.
- We propose an analysis that shows that the proposed quality indicators and the quality of the records have a mutual relationship, in which one depends on the other. We then show that indeed, outliers may represent inconsistent records with respect to the published literature.
- Using Principal component analysis (PCA) to reduce the dimensionality of data for visualization purposes, we show that records which have been reported as inconsistent with the literature fall roughly in the same area showing similar patterns.
- By manually analyzing records not previously known to be erroneous that fall in the same area than records know to be inconsistent, we show that 1 record out of 4 is inconsistent with respect to the literature (25% of the records are inconsistent in that area, denoted Zone B in Figure 4(a)). This high density of inconsistent record opens the way towards the development of automatic methods for the detection of faulty records.

The rest of this paper is organized as follows: in Section 2, we discuss the related work; in Section 3, we describe the list of quality indicators that we use to measure the literature consistency; in Section 4 we describe the experimental setup; in Section 5, we present the evaluation results and our analysis; and in Section 6, we conclude with key observations.

## 2 Related Work

There is a substantial body of research related to data quality in bioinformatics databases. Previous research has focused mainly on duplicate record detection and erroneous annotations, as reviewed below.

### 2.1 Duplicate records

Koh et al. [3] use association rule mining to check for duplicate records with per-field exact, edit distance, or BLAST sequence [8] alignment matching. Drawbacks of this method, and its poor performance, have been shown by Chen et al. [4]. Similarly, Apiletti et al. [9] proposed extraction of association rules among attribute values to find causality relationships among them. By analysing the support and confidence of each rule, the method can show the presence of erroneous data. Other approaches use approximate string matching to compute metadata similarity [10, 11, 12], under the assumption that duplicates have high metadata similarity.

Some approaches consider duplicates at the sequence level; they examine sequence similarity and use a similarity threshold to identify duplicates. For example, Holm and Sander [13] identified pairs of records with over 90% mutual sequence identity. Heuristics have been used in some of these methods to skip unnecessary pairwise comparisons, thus improving the efficiency. Li and Godzik [14] proposed CD-HIT, a fast sequence clustering method that uses heuristics to estimate the anticipated sequence identity and will skip the sequence alignment if the pair is expected to have low identity. Recently, Zorita et al. [15] proposed Star Code to detect duplicate sequences, which uses the edit distance as a threshold and will skip pairs exceeding the threshold. Such methods are valuable for this task, but do not address the problem of consistency or anomaly.

### 2.2 Erroneous annotations

Sequence databases exist as a resource for biomedicine, but the utility of the sequence of an organism depends on the quality of its annotations [12]. The annotations indicate the locations of genes and the coding regions in a sequence, and indicate what those genes do. That is, annotations serve as a reading guide to a sequence, which makes the scientific community highly reliant on this information. However, knowing that computational annotation methods introduce errors [5, 16, 17], the effect and the presence of these errors have been widely studied [18, 5, 17, 19, 1]. Some researchers then have developed strategies to detect and handle erroneous annotations. Weiser et al. [20] proposed Xanthippe, a system that is based on a simple exclusion mechanism and a decision tree approach using the C4.5 data-mining algorithm. Xanthippe uses the rules that are derivable from the learned decision tree to detect and flag a large proportion of the annotation errors; the method considerably increases the reliability of both automatically generated data and annotation from other sources.

Functional annotations are often inferred from sequence similarity to other annotated sequences, with the possibility of errors. The functional annotations in these sequences may themselves have been acquired through sequence similarity. Thus the possibility of erroneous annotation propagation may arise [17, 19]. Hence, Kaplan and Linial [6] proposed a protein-clustering method that enables automatic separation of annotations that are mistakenly associated with a protein from true hits. The method quantifies the biological similarity between pairs of proteins by examining each protein’s annotations, and then proceeds by clustering sets of proteins that received similar annotations into biological groups. Using a set of 327 test cases that are marked false positives, the authors show that their method successfully separates false positives in 69% of the cases.

Finally, Koh et al. [21] proposed a method for detection of attribute outliers from the correlation behavior of attributes. This method has been successfully applied to biological data, with results showing that this method is effective in identification of attribute outliers and detection of erroneous annotations.

Overall, existing data quality analysis methods for sequence databases focus only on the internal characteristics of records. Our work demonstrates that the literature associated to records is a valuable external source of information for assessing the quality of sequence database records.

## 3 What makes a sequence record consistent?

Our goal is to exploit the relationship between sequence records and the literature associated to those records. Hence, in this section, we first describe the structure of a sequence record, and then we describe a set of IR-based quality features.

### 3.1 Sequence record structure

The format of a sequence record can be regarded as having three parts [2]: the header, which contains the information that applies to the whole record; the features, which are the annotations on the sequence; and the sequence itself. The header section is composed of several fields: the *“LOCUS”* field, which contains a number of different data elements, including locus name, sequence length, molecule type, and modification date; the *“DEFINITION*” field, which is a brief description of sequence or sequence’s function; the *“ACCESSION NUMBER*”, which is a unique identifier for the record; the *“SOURCE*” field, which gives information about the sequence’s organism; and the *“REFERENCE*” field, which lists a set of publications by the submitters of the sequence that discuss the data reported in the record. It is clear that the header represents a rich source of information about the given sequence.

Knowing that the *“DEFINITION*” field of a record provides a succinct description of the sequence (information such as source organism, gene name/protein name, description of the sequence’s function, a completeness qualifier, etc.), we suggest to primarily use and focus on the information given in that field to assess the consistency of a record with respect to the literature.

### 3.2 Quality Factors

As stated previously, the *“DEFINITION*” field of a record provides a succinct description of the sequence, while research articles given in the *“REFERENCE*” field discuss the data reported in that record. Hence, adopting the approach of information retrieval (IR), if we consider the description given in the *“DEFINITION*” field of a record as a query into PubMed, the set of documents that can be considered as most relevant to that query are those listed in the “*REFERENCE*” field of that record. We propose to use IR methods to measure the quality of a record in terms of the quality of the query that is derived from the definition field of that record.

To find indicators or features to represent the quality of each query (record), we draw on the large body of previous work on query quality prediction [22, 23, 24]. These include predicting the quality of queries using either pre-retrieval indicators like Query Scope, that are calculated for a query as a whole, or post-retrieval indicators like Query Clarity, that involve assessing the results of an initial retrieval and hence are more expensive to compute. In the following, we describe the set of query quality predictors we used.

#### 3.2.1 Overlap similarity score

In IR a high quality query is a query that contains enough information (terms) to describe the user need. Hence, a high similarity value between a query *q* and its relevant documents *D* reflects a high query quality. We use the overlap similarity in order to emphasize the number of terms of a record definition that are in its set of associated articles. We use the maximum overlap similarity value obtained across the set of relevant documents to define this score. It is computed as follows:

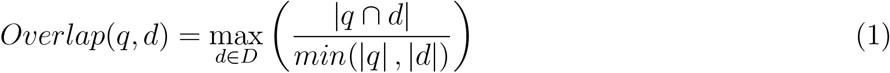

#### 3.2.2 Retrieval performance score (RP Score)

Based on the relevance paradigm of IR, we assume that a high quality query should retrieve and place its corresponding articles earlier in the ranking list. Thus, we use the Reciprocal Rank evaluation measure to define the RP Score as follows:

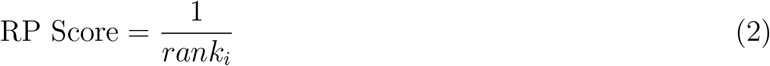

where *rank_i_* is the rank of the first relevant document in the retrieved list of documents that match the query *q* returned by the system. Note that we used the basic default Lucene vector-space model function to score and rank documents in order to compute the RP Scores.^8^

#### 3.2.3 Query scope (QS)

He and Ounis [25] defined Query Scope, an indication metric of the generality/speciality of a query based on the size of the document set containing at least one of the query terms. The basic idea is that a query that matches a high number of documents is likely to contain generic terms that don’t precisely express the user need (or respectively, the record definition contains generic terms that don’t precisely describe the sequence). Hence, we can expect that high values of query scope are predictive of poor-quality queries as they retrieve far too many documents. QS is computed as:

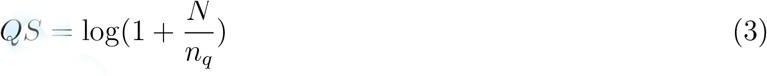

where *n_q_* is the number of documents that match the query terms, and *N* is the number of documents in the collection.

#### 3.2.4 TF score

The usage frequency of the query terms in the relevant documents is a good quality indicator of the query (respectively, the record’s quality). Clearly, a high overall term frequency means that the record is intensively discussed and reported in its associated articles. We propose to calculate a global value for the TF of the query terms across all relevant documents as follows:

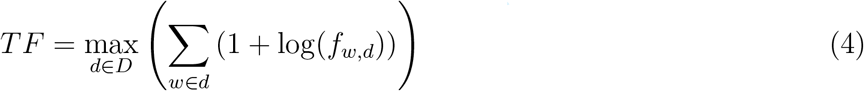

where *f_w_,_d_* is the number of time the term *w* occurs in the relevant document *d*.

#### 3.2.5 TF-IDF score

The TF-IDF (for Term Frequency-Inverse Document Frequency) is a term’s weighting function that estimates not only how important is a term to a document, but also considers how important is that term in the collection or the corpus. Hence, TF-IDF is defined such that a high value of is reached by a high term frequency and a low frequency of the term in the collection. We propose to calculate a global TF-IDF value for a query across all relevant documents as follows:

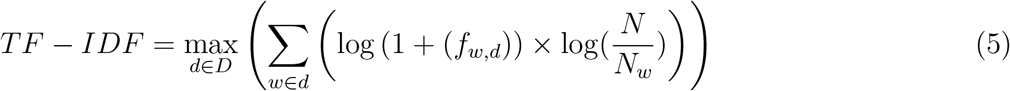

where *N_w_* is the document frequency of *w*. Like for the TF score, it is clear that a high global TF-IDF value indicates that the record is intensively discussed in the relevant documents while focusing on the main element reported in the record.

#### 3.2.6 Bayesian smoothing using Dirichlet priors score (LMDirichlet)

The approach of language modeling for information retrieval is based on defining a language model for each document *d*, and then estimating the conditional probability *p*(*d|q*), i.e., the probability that *d* generates the observed query *q*. Several variants of the language modeling technique for IR have been developed, most of them centered around the issue of smoothing. The term *smoothing* refers to the adjustment of the maximum likelihood estimator of a language model so that it will not assign a zero probability to unseen words. Zhai and Lafferty [26] studied the problem of language model smoothing and its influence on retrieval performance. They identified that a Dirichlet prior-based smoothing approach is among the most effective. The defined ranking function can be used to estimate the similarity between the record definition and the relevant documents. We refer the reader for the definition of this ranking function to the original article [26]. Note that in our previous work, this ranking function has been identified as the main discriminative function to identify dead records [7].

#### 3.2.7 Similarity of collection-query (SCQ)

Proposed by Zhao et al. [27], this query quality factor is based on the hypothesis that queries that have higher similarity to the collection as a whole will be of higher quality. SCQ is computed similarly to the TF-IDF weighting function, except that it focuses on the whole collection rather than on only the relevant documents. SCQ is computed as follows:

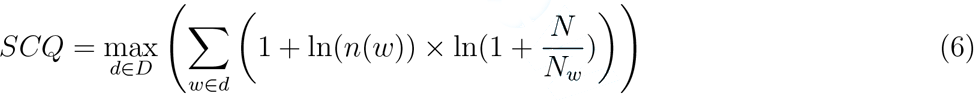

where *n*(*w*) is the frequency of the term *w* in the collection (sum of the frequencies of the documents in which *w* appears).

#### 3.2.8 Clarity score

The clarity score is post-retrieval query quality factor, which has been developed by Cronen-Townsend et al. [22]. It is simply the Kullback-Leibler divergence of the query model from the collection model. To avoid the expensive computation of query clarity, we computed the clarity score based on a query model estimated from the relevant documents themselves. We compute the clarity score as follows:

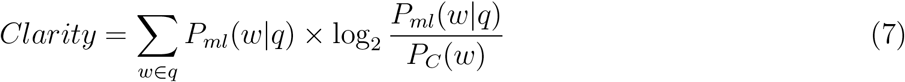

where *P_ml_*(*w|q*) is the probability of the occurrence of the word *w* in the relevant documents, and *P_C_*(*w*) is the probability of the occurrence of *w* in the collection. As stated by the authors of the clarity score, a query whose language model looks like the language model for the whole collection receives a low score, and a query whose language model is very different from the collection language model receives a high score. Therefore, a query with a high clarity value indicates the lack of ambiguity of that query, and thus, it indicates that this query is of a high quality.

In total, we have defined 8 IR-based quality features that leverage the literature associated to the records. All the quality factors defined above can be computed while separately considering each of the three different fields of the relevant documents namely, (i) the title, (ii) the abstract, and (iii) the body. This results in a total of 24 quality factors.

## 4 Data description

Now we provide details of the dataset that we evaluate in this paper.

### Articles

We used the PubMed Central Open Access collection^9^ (OA), which is a free full-text archive of biomedical and life sciences journal literature at the U.S. National Institutes of Health’s National Library of Medicine. The release of PMC OA we used contains roughly 1.13 million articles, which are provided in an XML format with specific fields corresponding to each section or subsection in the article. We used the Lucene IR System^10^ to index the collection, with the default settings for stemming and English stop-word removal. We defined a list of biomedical keywords which should not be stemmed or considered as stop-words, such as the protein names “THE” and “Is”. Each section of an article (title, abstract, body) is indexed separately, so that different sections can be used and queried separately to compute the quality features.

### Sequences

We work with the GenBank nucleotide database, but limit the sequence database records we work with to those that can be cross-referenced to the PMC OA article collection. Specifically, we used a regular expression to extract GenBank accession numbers mentioned in the PMC OA articles, thereby identifying literature that refers to at least one GenBank identifier. This resulted in a list of 733,779 putative accession numbers. Of these, 494,142 were valid GenBank nucleotide records that we were able to download using the e-utilities API [28]. Among the valid records, only 162,913 records also cite the corresponding articles, determined via title matching. This process gave us a list of 162,913 pairs of record accession numbers and PMC article identifiers, which cite each other.

Each record in this dataset was labelled as “alive” or “dead”, an attribute that we obtained using the eSummary API [28]. Note that the records that are reported as “dead” are explicitly labelled as such in GenBank, whereas records that are “alive” are implicitly labelled by not being labelled as dead. We consider dead records to be *suspicious* and all other records to be *confident* in our labelling.

In addition, “dead” records have been removed for several and different reasons, which are often given in a free text format. We grouped these removal reasons in the following categories:

1. Rm1: records were removed because the sequence was not directly determined by the submitter.
2. Rm2: records were removed at the submitter’s request because the sequence was determined to be incorrect.
3. Rm3: records were removed for unknown reasons.
4. Rm4: records were removed because no experimental evidence was published to support the records’ data.
5. Rm5: records were removed because the source organism cannot be confirmed.

Clearly, Rm4 represents the group of records that have been removed for literature inconsistency reasons. This group is the focus of our study, and hence, we have chosen to merge the other groups into a single group of removed records that we named “removed for other reasons”.

We provide more detailed statistics on our final dataset in Table 1. For example, an article cites 0.51 records on average; the article PMC2848993^11^ cites 8,062 records. A record is cited on average 1.17 times, while the record A23187^12^ is the most cited. Among the 162,913 records for which the relevant articles are in the PMC OA dataset, 162,486 are alive and only 427 are dead. Hence, our dataset is skewed toward many alive records and only very few dead records.

**Table 1:**
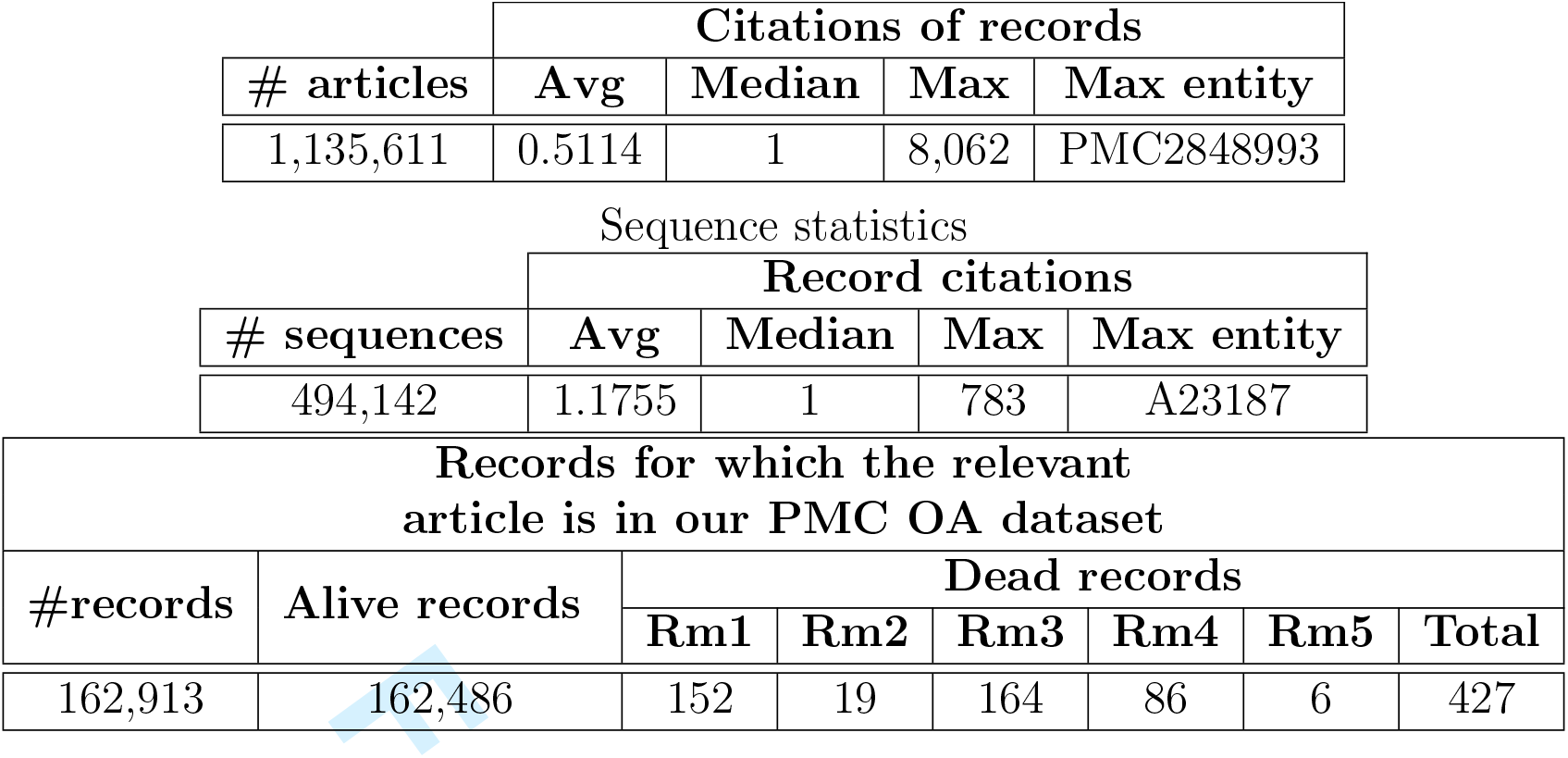
Dataset Statistics. Article statistics

## 5 Empirical evaluation

In this section, we first carry out a data quality feature analysis, then, we show and discuss the scatter plot obtained by applying Principal Component Analysis (PCA) for data visualization. Finally, we discuss the results obtained for the data curation task that we carried out.

### 5.1 Quality Features Analysis

We analyze the literature consistency of our records using the metrics defined in Section 3.2, To do so, we compare the distribution of the quality scores obtained for two groups of records: (i) Rm4, the group of removed records because of literature consistency reasons, and (ii) the group of alive records. We visualize the analysis using a violin-plot, which is a similar data visualization method to box-plot except that it also shows the probability density of the data at different values on each side. We also include a marker for the median and the inter-quartile range of the data that we use for comparison.

Figure 2 shows grouped and split violin-plots of the quality scores for both alive and records of Rm4 group. We also show a violin-plot for each metric computed based on different sections of the articles (title, abstract, and body). At first glance, it is clear that for all metrics, the distribution of values for the two groups are somehow different.

**Figure 2:**
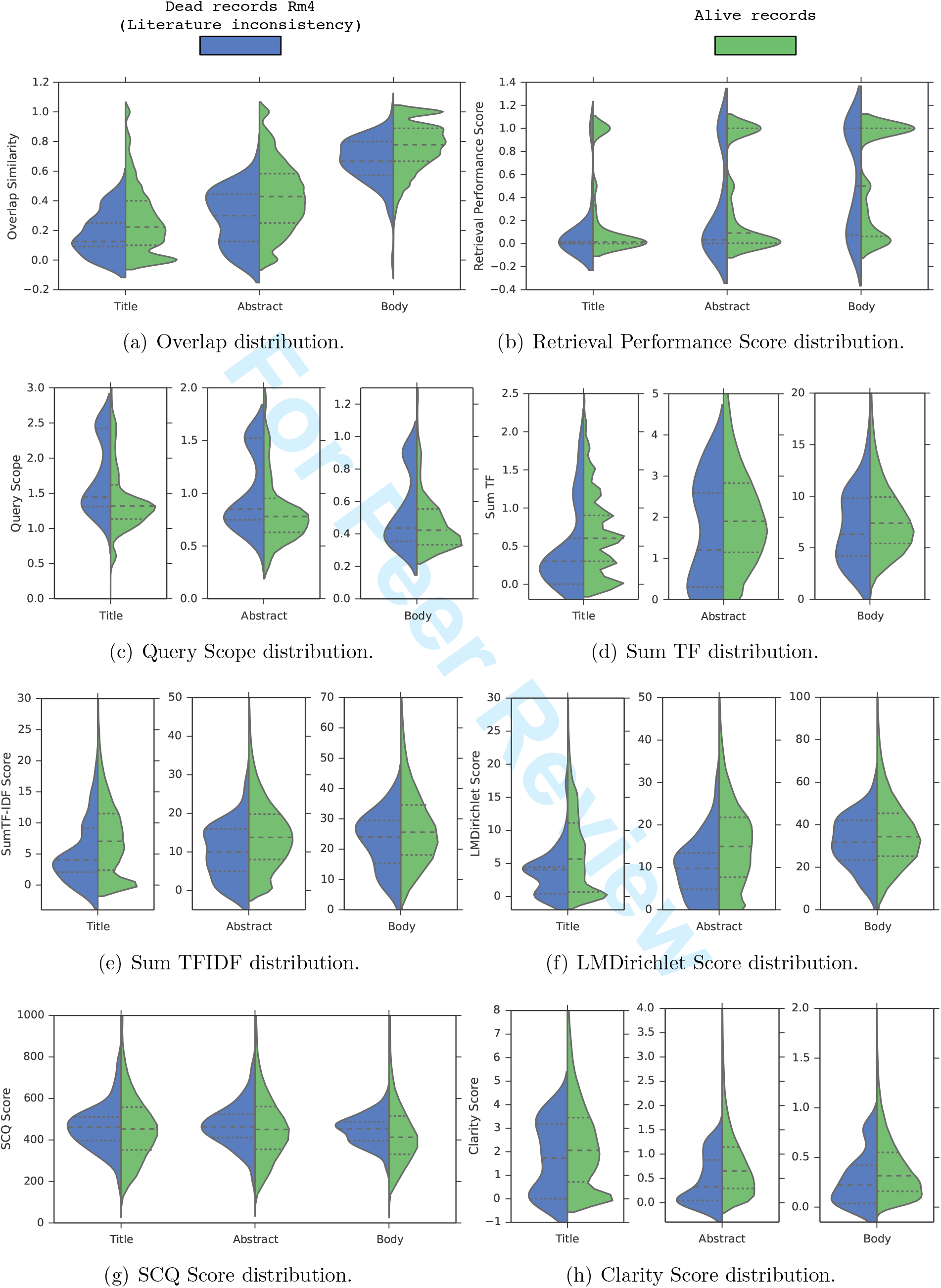
Grouped and splitted violin-plots of the features for both alive and dead records.

First, regarding the overlap similarity distribution shown in Figure 2(a), we notice four notable trends: first, term overlap increases from title to body and full text since the size grows accordingly; second, overall, for alive records, there is a high term overlap of roughly 80% between the descriptions and the body sections; and third, for a small number of records, in which the overlap similarity is below 0.2, there is low overlap or no overlap at all between the description field and the full text of their associated articles, thus suggesting a data quality problem; and last but not least, Figure 2(a) shows clearly that literature-inconsistent records (Rm4 group) tend to have a lower overlap similarity across all articles parts compared to the alive records as shown by the median lines. Especially, for the three compared distributions, the third quartile of the Rm4 group is almost always aligned with the median of the alive records showing that 50% of the alive records tend to have a higher overlap similarity than 75% of the records in the Rm4 group. Hence, we conclude that the overlap similarity is a strong indicator for the literature consistency of the records.

As for the distribution of Retrieval Performance (RP) scores illustrated in Figure 2(b), the difference is unclear. Indeed, the RP score distributions using the title and the body are similar for the two groups since the quartiles and the medians are aligned. However, the RP score distribution for the abstract is slightly different since the medians and the third quartiles are not aligned. The alive records tend to rank higher the relevant documents than the Rm4 group when querying the abstract sections. This shows that the initial assumption of this quality score is somehow verified.

Figure 2(c) shows the distribution of the QS scores. For the three article parts, it is clear that in general, the records in the Rm4 group tend to have higher QS scores than the alive records. This observation validates the intuition behind the QS score which is that inconsistent records are likely to contain generic terms that don’t precisely describe the sequences. In particular, we notice that the first quartile of the Rm4 group is almost aligned with the median of the alive records when querying the titles or abstracts. This again shows that 50% of the alive records tend to contain less generic description terms than 75% of the records in the Rm4 group. Hence, we conclude that the QS metric is a good indicator for the literature consistency of the records.

Figures 2(d) and 2(e) show the distributions of the TF and TFIDF scores respectively. Again, overall, for the two metrics, and for the three article parts, the records in the Rm4 group tend to have a lower TF and TFIDF scores than the alive records. This indicates that overall, the records in the

Rm4 group are less discussed and reported in their articles than the alive records, which is a literature inconsistency indicator.

Also, Figure 2(f) shows the distribution of the LMDirichlet scores. Again, for the three articles parts, it is clear that roughly, the records in the Rm4 group tend to have lower LMDirichlet scores than the alive records. This indicates that literature-inconsistent records are less likely to have been generated from the articles. In particular, we notice that the third quartile of the Rm4 group is almost aligned with the median of the alive records when using the titles or abstracts. This again shows that 50% of the alive records tend to be likely to have been generated from their research articles than 75% of the records in the Rm4 group. Hence, we also conclude that the LMDirichlet metric is a good indicator for the literature consistency of the records.

Figure 2(g) shows the distribution of the SCQ scores. Here, the difference is unclear, showing that the distributions are similar between the records of the Rm4 group and the alive records. This analysis shows clearly that SCQ is not a good quality factor to detect records inconsistent with the literature.

Finally, Figure 2(h) shows the distribution of the clarity scores. Roughly speaking, the distributions show that the alive record tend to have higher clarity scores than the records in the Rm4 group.

This indicates that in general, the definition of the records in the Rm4 group are less clear than the alive records. This verifies the assumption behind the clarity score in the context of the detection of literature-inconsistent records in bioinformatics sequence databases.

In summary, this quality feature analysis shows that, except the SCQ and the RP metrics, all the quality factors described in Section 3.2 reveal patterns that confirm our intuitions in the context of the detection of literature-inconsistent records. This opens new avenues to develop automatic/semi-automatic methods to assist curators in detecting inconsistent records to ensure a high database quality.In the next section, we carry out a data visualization/exploration task.

### 5.2 PCA for data visualization

In many Machine Learning problems, visualizing the data provides insight into its solvability. Hence, several dimensionality reduction methods have been developed for that purpose.

In our dataset each record *m* is represented by its vector of 24 quality indicators 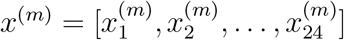(see Section 3.2). Visualizing this 24-dimensional data is very difficult. Using dimensionality reduction, we can reduce these 24D into a 2D data, and visualizing 2D data is much simpler. However, the dimensionality reduction algorithm doesn’t give any meaning to the new features; it is up to us to figure out what it means. Principal Component Analysis (PCA) is the most commonly used dimensionality reduction algorithm. PCA finds a vector direction onto which the data are projected and which gives the smallest sum of squared projected errors. We refer the reader to the book of Jolliffe [29] for more details about PCA.

In this paper, we use PCA for a 2D visualization of our dataset as depicted in Figures 3(a) and 3(b), with 46.41% of variance retained in the two first components. Both figures show a scatter plot of the alive records with a Gaussian kernel density representation. The red points represent the centroid of the alive records, whereas the blue points may be considered as outliers. In addition, each figure shows also the dead records using orange points, i.e., the records removed for inconsistency reasons in Figure 3(a), and those removed for other reasons in Figure 3(b). The plot in Figure 3 leads us to make the following two observations: [label=(v)]

1. First, in Figure 3(b) the orange points representing records removed for reasons other than literature inconsistency have the same probability density as the alive records. This indicates that, using the quality factors defined in Section 3.2, it is not possible to detect records that have been removed for other reasons not related to literature consistency, such as sequence correctness.
2. Second, in Figure 3(a) the orange points representing records removed for literature inconsistency have a different probability density than the alive records. It appears that these records tend to be more outliers, falling in the south region of the scatter plot.

These two observations led us to ask several research questions:

- What do alive outlier records represent (blue points)?
- Do all alive outlier records contain anomalies related to literature inconsistency? Or in contrast, are they highly consistent with the literature?
- Can we find inconsistent records among the alive ones within the red points (the centroid region)?
- Can we define an automatic/semi-automatic method to detect inconsistent records based on our quality metrics?
- How accurate is a clustering based method to detect literature-inconsistent records? Can we achieve better performance than a random selection approach?

To answer all the questions above, we carried out a data curation task that we describe in the next section.

**Figure 3:**
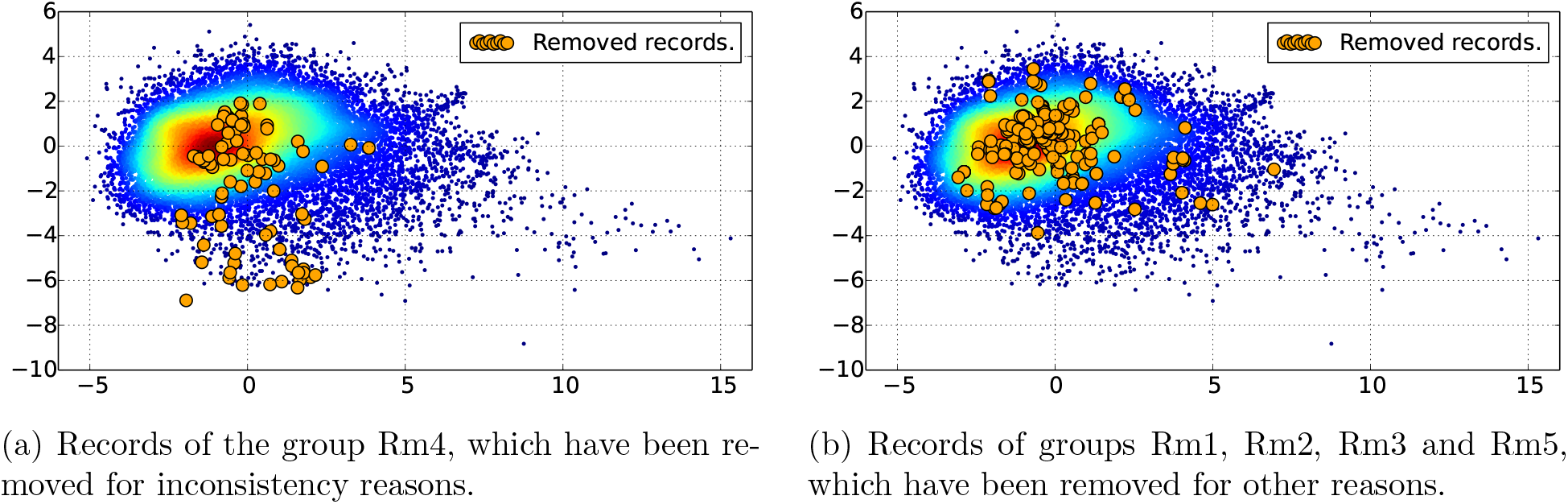
Dataset plotted in 2D, using PCA for dimensionality reduction with 46.41% of variance retained in the two first components.

### 5.3 Data curation task

The main objective of this data curation task is to explore literature-based consistency assessment. To address the questions raised in the previous section, we have built a data curation test set, based on the scatter plot of Figure 3(a). We divided the data into three interesting zones to explore, as illustrated in Figure 4(a). Zone A covers the centroid region, Zone B covers the south outlier region, and Zone C covers the east outlier region. Figure 4(b) shows the records effectively selected to be included in the test dataset for the curation task. The zones B and C contain 100 records each. In region A, which is very dense, we have randomly selected 100 records. Additionally, we have randomly selected 100 records from the whole dataset to be included in the test set for comparison. This gave us a total of 400 records to be curated in order to answer the questions previously posed.

**Figure 4:**
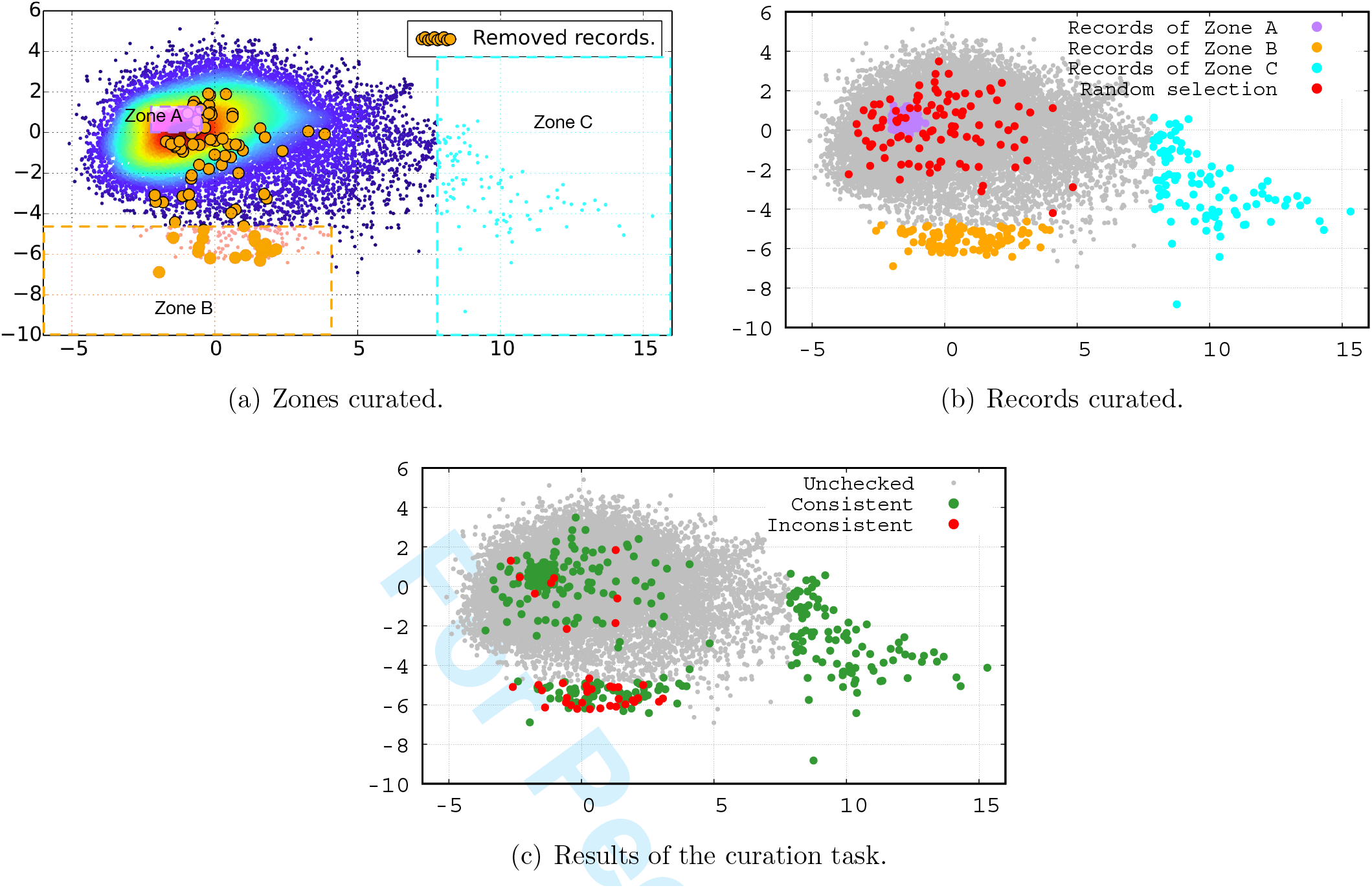
Curation task

We hired a University of Melbourne Masters student in Bioinformatics with an undergraduate degree in Biology to perform the job of a database curator. For each record to be assessed, we have given the student the full content of the record and access to each article listed by that record. The student was then asked to check the record content, and to review each article to look for inconsistencies. Inconsistent records include records which either (i) cite the wrong organism, i.e., a different organism than the one discussed in its associated articles, (ii) cite a different gene name (or protein name) than the one discussed in its associated articles, or (iii) focus on a gene (or a protein) that is not reported in its associated articles. However, we were interested in any form of inconsistency, and thus, the student was asked to freely consider inconsistency cases other than those listed here.

The student was trained using a separate training dataset of 30 records, among which 10 were known to be inconsistent with the literature, and 20 were consistent. The 30 records were incrementally presented to the student in three groups of ten records. The student was then asked to indicate for each record whether she judged the record as consistent with its associated articles, or not. The student was able to gradually improve her performance on the task until she achieved a 100% accuracy on the third group. We then asked her to check the consistency of the records of the test set. The results of the curation task are illustrated in Figure 4(c) for visualization purposes, and provided with more details in Table 2.

**Table 2:**
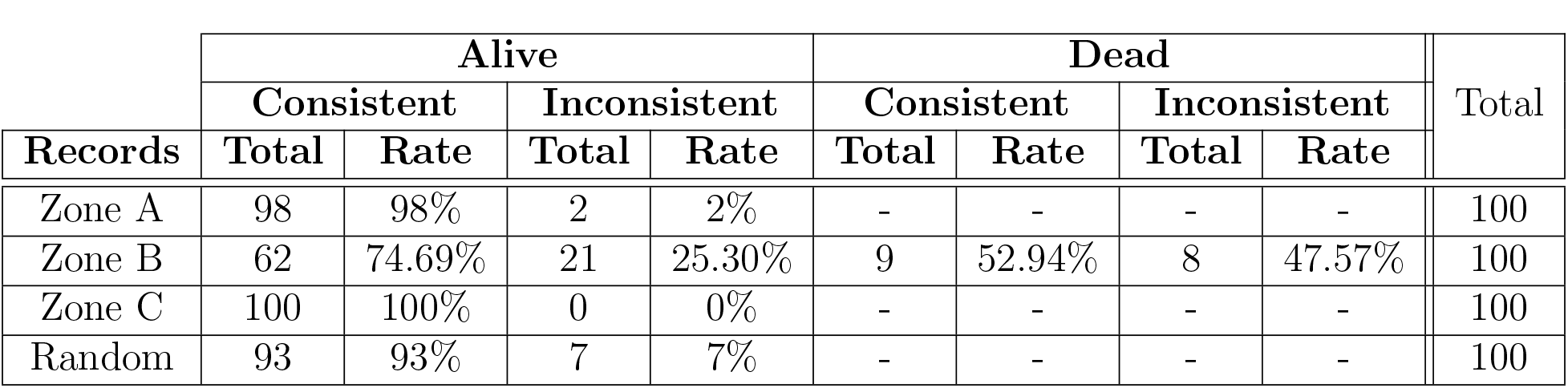
Details of the results of the curation task.

| Records | Alive |  |  |  | Dead |  |  |  | Total |
| --- | --- | --- | --- | --- | --- | --- | --- | --- | --- |
|  | Consistent |  | Inconsistent |  | Consistent |  | Inconsistent |  |  |
|  | Total | Rate | Total | Rate | Total | Rate | Total | Rate |  |
| Zone A | 98 | 98% | 2 | 2% | - | - | - | - | 100 |
| Zone B | 62 | 74.69% | 21 | 25.30% | 9 | 52.94% | 8 | 47.57% | 100 |
| Zone C | 100 | 100% | 0 | 0% | - | - | - | - | 100 |
| Random | 93 | 93% | 7 | 7% | - | - | - | - | 100 |

The main outcome of this curation task can be summarized as follows:

i. The records of Zone C were all identified as being highly consistent. They present similar profiles of a complete genome of a given organism, for which a research article is assigned in order to describe this genome in detail.
ii. In Zone A, 98% of the records were marked as consistent by the student. Only two records have been identified as inconsistent. The record with the accession number KF188702^13^ cites a different isolate^14^ code than the one cited in its associated article. The other record with the accession number KC164376^15^ was reported as inconsistent with the following comment *“not too sure about what the species is in the record, record says ‘synthetic construct’ but it’s not mentioned at all in the article”*.
iii. The random selection methods allowed to identify up to 7% of inconsistent records.
iv. In Zone B, up to 25% of the records have been identified as being inconsistent. In other words, in Zone B, 1 record out of 4 is inconsistent with respect to the literature. This confirms that literature-inconsistent records have a similar profile, which therefore should lend themselves to identification using machine learning methods. We infer from this result that automatic methods can be developed to help curators in assessing only records that have a high probability of being inconsistent.
v. Finally, Zone B contains 17 records which are currently reported as being inconsistent, and thus are dead. It is interesting to notice that 8 of these records have been reported as being consistent by our curator. We explain that by the fact that our curator may have insufficient knowledge to detect latent inconsistencies (not obvious or explicit inconsistencies). Indeed, our curator acknowledged and mentioned the difficulty in judging the consistency of several records, mostly because of the broad knowledge needed to assess them. Those records for which the consistency was difficult to reject were marked as consistent by our curator. This suggests that the current data quality status is worse than one may think, in that there are certainly records that have been judged as consistent in the “Alive” data that are in fact inconsistent based on characteristics that require deeper expertise.

## 6 Conclusions and future work

In this paper, we have proposed an analysis of data quality in bioinformatics sequence databases from the perspective of consistency with respect to the published research literature. We have introduced a list of factors that have a mutual dependence with the consistency of a record with respect to its associated articles. We then compared the distribution of the values of each quality factor for records which have been reported as inconsistent with the literature, with those currently alive in our dataset.

Using PCA to reduce the dimensionality of data for visualization purposes, we showed that records which have been reported as inconsistent with the literature fall roughly in the same area, showing similar patterns according to our proposed quality factors. By manually analyzing the “alive” records that fall in the same area than records know to be inconsistent, we show that 1 record out of 4 is inconsistent with respect to the literature (25% of the records are inconsistent in that area, denoted Zone B in Figure 4(a)). This high density of inconsistent record opens the way towards the development of clustering methods for the detection of faulty records. The main outcome of this work is therefore that literature inconsistency shows promise for helping to flag suspicious sequence records.

This work is to the best of our knowledge the first that focuses on identifying inconsistent records in biomedical sequence databases with respect to the published literature. We have shown that automatic or semi-automatic methods can be developed to help curators in identifying such anomalies in order to potentially greatly reduce the effort required by curators to identify and remove low-quality records.

Future work includes the use of methods such as k-nearest neighbor or local outlier factor [30] to detect inconsistent records. We believe that we can achieve high precision/recall scores using a machine learning classifier devoted to the detection of outliers. We also envision to release a benchmark data set for this task, which we hope would attract much attention from the biocuration community in the future.

## Acknowledgement

We would like to thank Keely Aranyos for her work as a database curator.

## Footnotes

1 http://www.ncbi.nlm.nih.gov/genbank

2 http://www.ncbi.nlm.nih.gov/genbank/statistics/ (release 217)

3 http://www.ncbi.nlm.nih.gov/nuccore/BK004887.1?report=genbank

4 http://www.ncbi.nlm.nih.gov/pmc/articles/PMC555591/

5 http://www.ncbi.nlm.nih.gov/nuccore/FJ824848

6 A cloning vector is a small piece of DNA, taken from a virus, a plasmid, or the cell of a higher organism, that can be stably maintained in an organism, and into which a foreign DNA fragment can be inserted for cloning purposes.

7 http://www.ncbi.nlm.nih.gov/pmc/articles/PMC2675058/

8 https://lucene.apache.org/core/6_2_0/core/org/apache/lucene/search/similarities/TFIDFSimilarity.html

9 http://www.ncbi.nlm.nih.gov/pmc/tools/openftlist/

10 http://lucene.apache.org/

11 http://www.ncbi.nlm.nih.gov/pmc/articles/PMC2848993/

12 http://www.ncbi.nlm.nih.gov/nuccore/A23187

13 http://www.ncbi.nlm.nih.gov/nuccore/KF188702

14 “Isolate” refers to the place where the organism has been sampled.

15 http://www.ncbi.nlm.nih.gov/nuccore/KC164376

